# Distribution-based measures of tumor heterogeneity are sensitive to mutation calling and lack strong clinical predictive power

**DOI:** 10.1101/248435

**Authors:** Javad Noorbakhsh, Hyunsoo Kim, Sandeep Namburi, Jeffrey Chuang

## Abstract

Mutant allele frequency distributions in cancer samples have been used to estimate intratumoral heterogeneity and its implications for patient survival. However, mutation calls are sensitive to the calling algorithm. It remains unknown whether the relationship of heterogeneity and clinical outcome is robust to these variations. To resolve this question, we studied the robustness of allele frequency distributions to the mutation callers MuTect, SomaticSniper, and VarScan in 4722 cancer samples from The Cancer Genome Atlas. We observed discrepancies among the results, particularly a pronounced difference between allele frequency distributions called by VarScan and SomaticSniper. Survival analysis showed little robust predictive power for heterogeneity as measured by Mutant-Allele Tumor Heterogeneity (MATH) score, with the exception of uterine corpus endometrial carcinoma. However, we found that variations in mutant allele frequencies were mediated by variations in copy number. Our results indicate that the clinical predictions associated with MATH score are primarily caused by copy number aberrations that alter mutant allele frequencies. Finally, we present a mathematical model of linear tumor evolution demonstrating why MATH score is insufficient for distinguishing different scenarios of tumor growth. Our findings elucidate the importance of allele frequency distributions as a measure for tumor heterogeneity and their prognostic role.

## BACKGROUND

A major challenge for predicting clinical outcome to cancer treatment is the heterogeneity of cell populations within each tumor, as intratumoral heterogeneity is increasingly being associated with metastasis and resistance to therapies [1–4]. Intratumoral heterogeneity develops from mutations in cells and the relative growth advantages they confer to their descendant populations [5, 6]. To better understand this phenomenon, multiple research groups have developed methods for the challenging problem of estimating tumor cellular composition from bulk tumor sequencing data [7–10]. These methods rely on the allele frequencies of mutations in each cancer, with low allele frequency mutations providing information on rare subclones and high allele frequency mutations providing information on common subclones. Although factors such as ploidy, copy number, and tumor purity can affect this inference, allele frequencies are essential for evaluating subclonal heterogeneity and its clinical implications.

Increased heterogeneity has been theorized to lead to worse patient survival due to the increased potential for resistant populations [11–13], but it remains unclear if current measures of tumor heterogeneity are sufficient to resolve such an effect. Prior studies have attempted to determine this relationship. For example, Rocco et. al. [10] used MATH score, a measure of the width of the allele frequency distribution, as a proxy for tumor heterogeneity and observed poorer survival for head and neck squamous cell carcinoma (HNSC) in patients with higher MATH score. More recently, Morris *et. al.* [14] found associations between heterogeneity and survival in several cancers using a multivariate regression of MATH and other variables versus survival. However, a key caveat is that all heterogeneity measures, including MATH, are affected by the accuracy of mutation calls. Many studies have shown that cancer mutation calls can differ substantially depending on the algorithm used for their determination [15–18]. In order to verify if heterogeneity impacts survival, it is necessary to quantify the robustness of allele frequency distributions to mutation callers as well as the robustness of their relationship to survival.

Allele frequency distributions also provide information on the evolutionary processes in tumors, which remain poorly understood. While a variety of intratumoral evolutionary models have been proposed [19–22], the impact of allele frequency accuracy on evolutionary inference has not been substantially explored. Determining the robustness of allele frequency distributions will elucidate this problem.

In this paper we study the robustness of allele frequency distributions and their relevance to patient survival. To the best of our knowledge this is the first study which explores sensitivity of these distributions and their clinical prognostic power to different mutation calling methods. We call mutations from 11 cancer types in The Cancer Genome Atlas (TCGA) using three common mutation callers: MuTect [23], SomaticSniper [24], and VarScan [25, 26]. To determine if the resulting allele frequency distributions are clinically useful, we analyze the correlation between these distributions and patient survival. Our study demonstrates whether allele frequency variability is clinically predictive and what other genomic features mediate these results. Finally, we discuss implications for evolutionary mechanisms of resistance.

## RESULTS

### Different mutation callers lead to distinct allele frequency distributions

We analyzed a total of 4722 samples from the TCGA database on a cloud computing platform to explore the effects of mutation calling by MuTect, SomaticSniper, and VarScan on allele frequency distributions and patient survival. Tumor/normal matched BAM files that had been aligned to the hg19 reference were used to call the somatic mutations (**Figure 1A**). The mutation calling pipelines (**Figure S1**) were implemented by dockerizing [27] each element of the pipeline and linking them through an interface provided by the Cancer Genome Cloud (CGC) [28]. Three different mutation calling pipelines were run for each tumor/normal pair on Amazon Web Services (AWS) [29] through the CGC interface, and the resulting allele frequencies were calculated for each sample.

**FIG 1.**
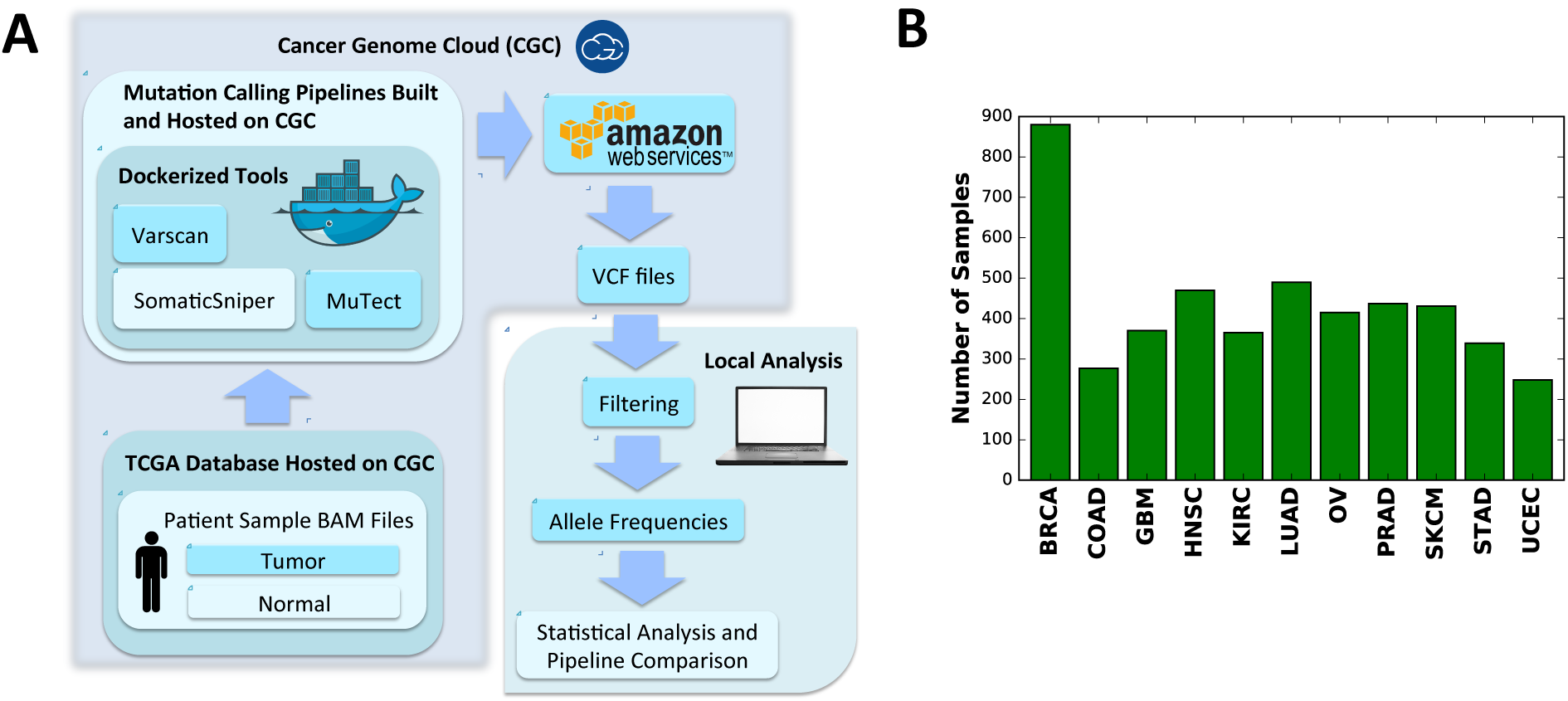
Analysis design. A) Schematic of the analysis process for 4722 TCGA samples. Somatic mutation calling was carried out on the NCI Cancer Genomics Cloud [28] with Amazon Web Services [29] backend, followed by additional local analysis. B) Number of samples studied from each cancer.

To analyze the range of possible allele frequency behaviors, we chose 11 cancer types from the TCGA database (**Figure 1B**), namely breast invasive carcinoma (BRCA), colon adenocarcinoma (COAD), glioblastoma multiforme (GBM), head and neck squamous cell carcinoma (HNSC), kidney renal clear cell carcinoma (KIRC), lung adenocarcinoma (LUAD), ovarian serous cystadenocarcinoma (OV), prostate adenocarcinoma (PRAD), skin cutaneous melanoma (SKCM), stomach adenocarcinoma (STAD), and uterine corpus endometrial carcinoma (UCEC).

We then compared allele frequency distributions assessed by pairs of mutation callers using the Kolmogorov-Smirnov test (**Figure 2A,B**). Distributions were not very robust across different cancer types (**Figure 2B**). SomaticSniper and VarScan produced the most dissimilar allele frequency distributions with 42 ± 14% (mean±std) of samples being significantly different across cancers. MuTect and VarScan produced the least dissimilar results with 11 ± 8% of samples significantly different. This percentage was 22 ± 14% when MuTect was compared against SomaticSniper. Prostate adenocarcinoma (PRAD), showed unusually robust results, with only 1%, 1%, and 3% of samples being significantly dissimilar when MuTect-SomaticSniper, MuTect-VarScan, and SomaticSniper-VarScan were compared pairwise, respectively.

**FIG 2.**
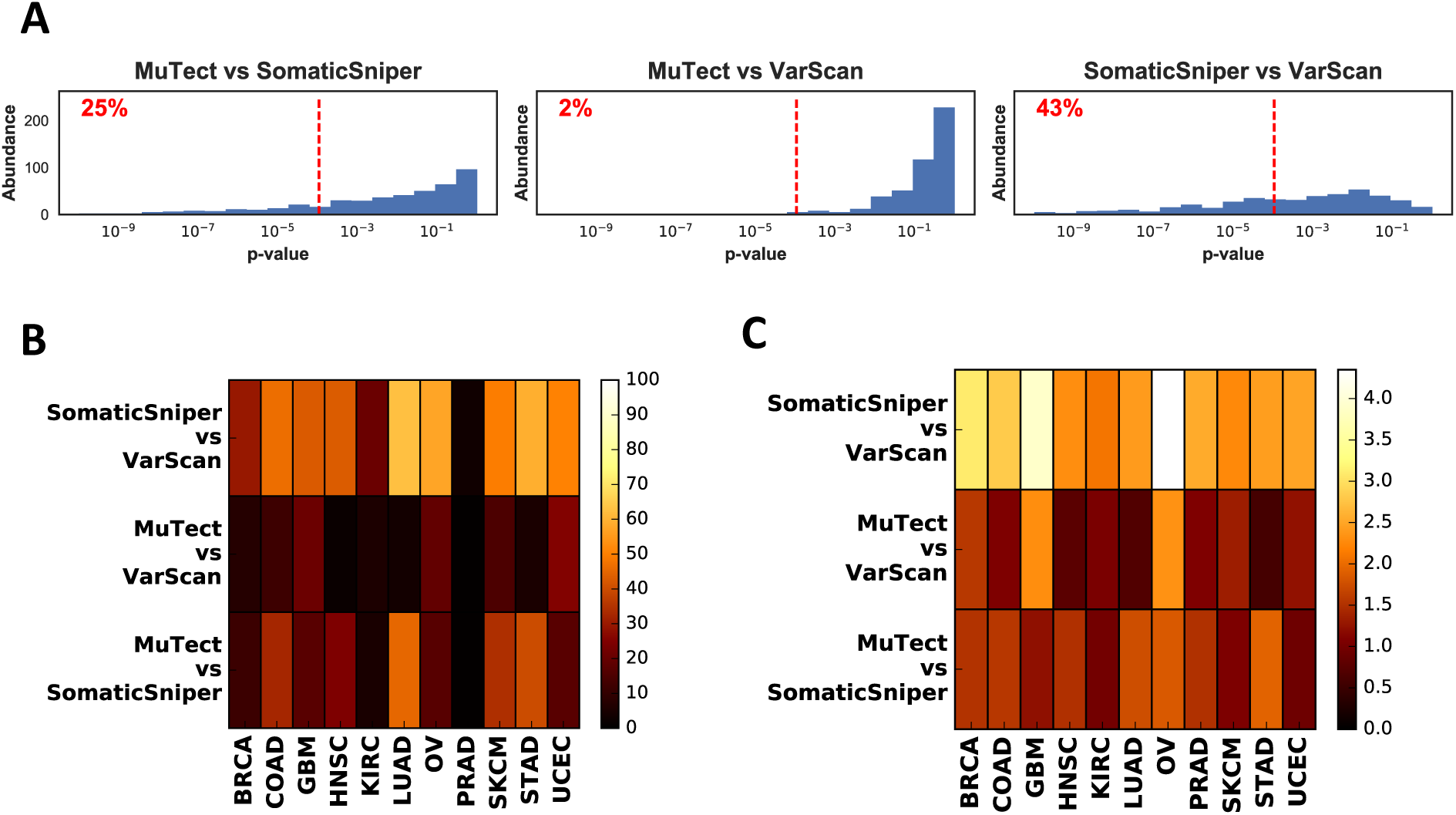
Comparisons of allele frequency distributions for different mutation callers. A) Distribution of Kolmogorov-Smirnov (KS) test p-values corresponding to the null hypothesis that mutation callers will produce similar allele frequency distributions, plotted pairwise for HNSC. The red dashed line indicates the Bonferroni corrected significance threshold *α* = 0.05. The percentage of samples that fall below this threshold are shown on each graph. B) Percentage of significantly different samples for mutation caller pairs, grouped by cancer type. C) Comparison of allele frequency distributions using earth mover’s distance.

Due to sensitivity of the Kolmogorov-Smirnov test to the number of mutations in the tumor (**Figure S2**), we also used the earth movers distance (EMD) to assess differences between allele frequency distributions. EMD produced qualitatively similar results (**Figure 2C**) to the Kolmogorov-Smirnov test, with SomaticSniper-VarScan showing the most difference among all pairwise comparisons and MuTect-VarScan showing the least difference. A third method based on cumulative absolute differences between pairs of distributions also yielded similar results (**Figure S3**, and *Methods*).

Copy number variations (CNV) can influence allele frequencies and may indirectly shape their distribution. To assess this effect, we repeated the analysis after removing somatic mutations with copy number aberrations (|*CNV*| > 0.2). Comparison of allele frequency distributions using Kolmogorov-Smirnov statistics (**Figure S4**) did not produce qualitatively different results from those shown in **Figure 2B**, and again showed the most pan-cancer differences between SomaticSniper and VarScan. These results suggest that copy number impacts all mutation callers similarly.

### MATH score is a poor predictor of patient survival across cancer types

To investigate the relationship of tumor heterogeneity and survival, Mroz *et al.* [10] introduced a measure of heterogeneity they term MATH. It is the ratio of scaled median absolute deviation (MAD) to median stated in percentage:

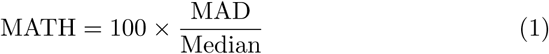

We calculated this measure for all samples and mutation callers. Overall VarScan yielded higher median MATH score (35.4±5.7) compared to SomaticSniper (24.4±4.8) and MuTect (30.3 ± 5.9) (**Figure S5A**). Similar to **Figure 2**, the MATH scores were also more similar between MuTect and SomaticSniper calls (pan-cancer Spearman correlation coefficient *ρ* = 0.7 ± 0.1) and more dissimilar between SomaticSniper and VarScan (*ρ* = 0.5 ± 0.2) (**Figure S5B**).

We then analyzed the relationship between MATH score and patient survival by grouping samples into high or low MATH groups as compared to the median of the cohort. These two groups were then compared using a log-rank test to determine the significance of MATH values on Kaplain-Meier patient survival curves (**Figure 3** and **Figure S6**).

**FIG 3.**
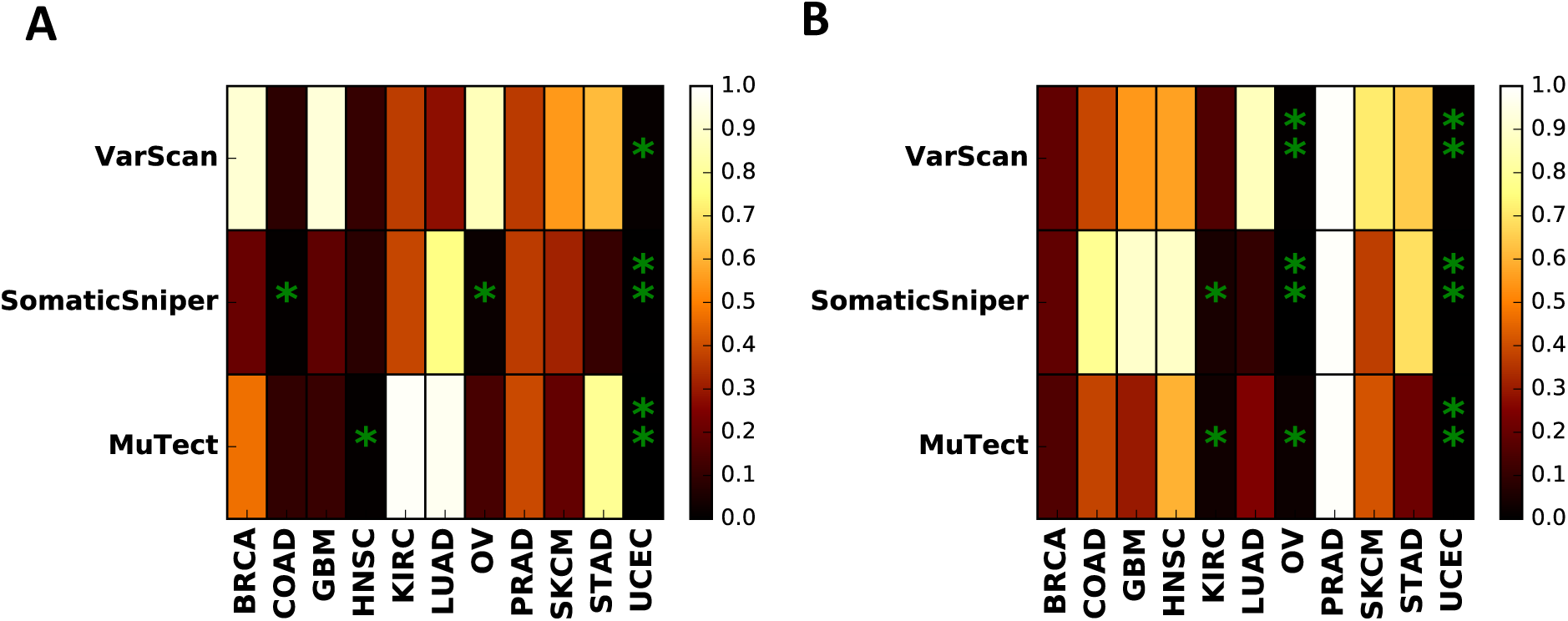
Significance analysis of measures of genomic variations and patient survival. A) Survival analysis log rank test p-values for high and low MATH score groups, and B) Survival analysis log rank test p-values for high and low CNV standard deviation groups. Stars correponds to values smaller than significance threshold 0.05. Double stars show significant results after Benjamini-Hochberg correction across all cancer types and mutation callers.

Among the cancers studied, COAD, OV, HNSC, and UCEC showed a significant (*p* < 0.05) relationship between MATH and survival for at least one caller (e.g. COAD p=.007, OV p=0.02 with SomaticSniper calls). However, only UCEC showed a significant relationship for all three mutation callers (p=.00083, 0.0023, .011 for MuTect, SomaticSniper, and VarScan calls, respectively). Although these results suggest some clinical predictive power of MATH score, a more conservative approach would be to correct for multiple hypothesis testing. Using the Benjamini-Hochberg correction for all cancers and mutation callers led to significant results only for UCEC, and only when the calls were made by MuTect or SomaticSniper. Therefore the predictive power of MATH score is not robust in a pan-cancer analysis except for possibly UCEC.

Previous studies [10, 14] have shown a significant relationship between MATH score and patient survival in HNSC. We obtained similar results when the calls were made using MuTect; however the effect was not significant when other callers were used (**Figure S6**). Restricting the analysis to the same samples as in [14] did not affect the significance of this result (**Figure S7**). Thus the clinical predictive power of MATH in HNSC appears to be substantially impacted by the mutation calling method.

To determine the direction of effect of MATH score on survival, we calculated the average survival difference between high and low MATH score groups (see *Methods*). This analysis indicates that for UCEC higher MATH score is associated with poorer patient survival (**Figure S8A**). The same trend is observed for COAD and HNSC. For OV, the direction of the trend is caller-dependent.

### Copy number mediates prognostic power of allele frequency distributions

Allele frequency variation in a tumor is impacted by locus-to-locus copy number variation in addition to subclonal heterogeneity, and decomposing these two contributions may be important for clinical prognosis. We therefore repeated the above survival analysis but replaced MATH score with the standard deviation of CNV over mutated loci (**Figure 3B** and **Figure S9**). This CNV analysis yielded more robust results across mutation callers than the MATH score analysis (**Figure S10**). Specifically, we found that copy number variation is correlated with survival in KIRC, OV, and UCEC for at least two mutation callers. Multiple hypothesis correction led to a robust effect in UCEC for all mutation callers as well as significance in OV for SomaticSniper and VarScan. This result indicates that copy number variation is a more robust predictor of patient survival than allele frequency distribution.

Despite the lack of statistical significance, for most cancers, higher CNV standard deviation led to poorer survival and this effect was robust across mutation callers (**Figure S8B**). This was most pronounced for SKCM, STAD, and UCEC. An exception was OV where higher copy number standard deviation was predictive of better survival.

To further distinguish the effect of allele frequencies and copy numbers on patient survival, we filtered out mutant allele frequencies at loci with |*CNV*| > 0.2 and calculated the log rank test for MATH using the remaining sites (**Figure S11**). Analysis of this set with multiple hypothesis correction did not yield any significant results, indicating that MATH clinical predictions are indeed driven by copy number aberrations. This result can be more clearly seen from the correlation between allele frequency variation and copy number variation. Pearson correlation coefficients between copy number standard deviation and MATH score were positive for all tumor types and all mutation callers (**Figure 4A**). This effect was even stronger when we compared copy number standard deviation to allele frequency standard deviation (**Figure 4B**). This further supports that allele frequency distributions are highly influenced by copy number aberrations across the genome (see examples in **Figure 4C** and **Figure 4D**).

**FIG 4.**
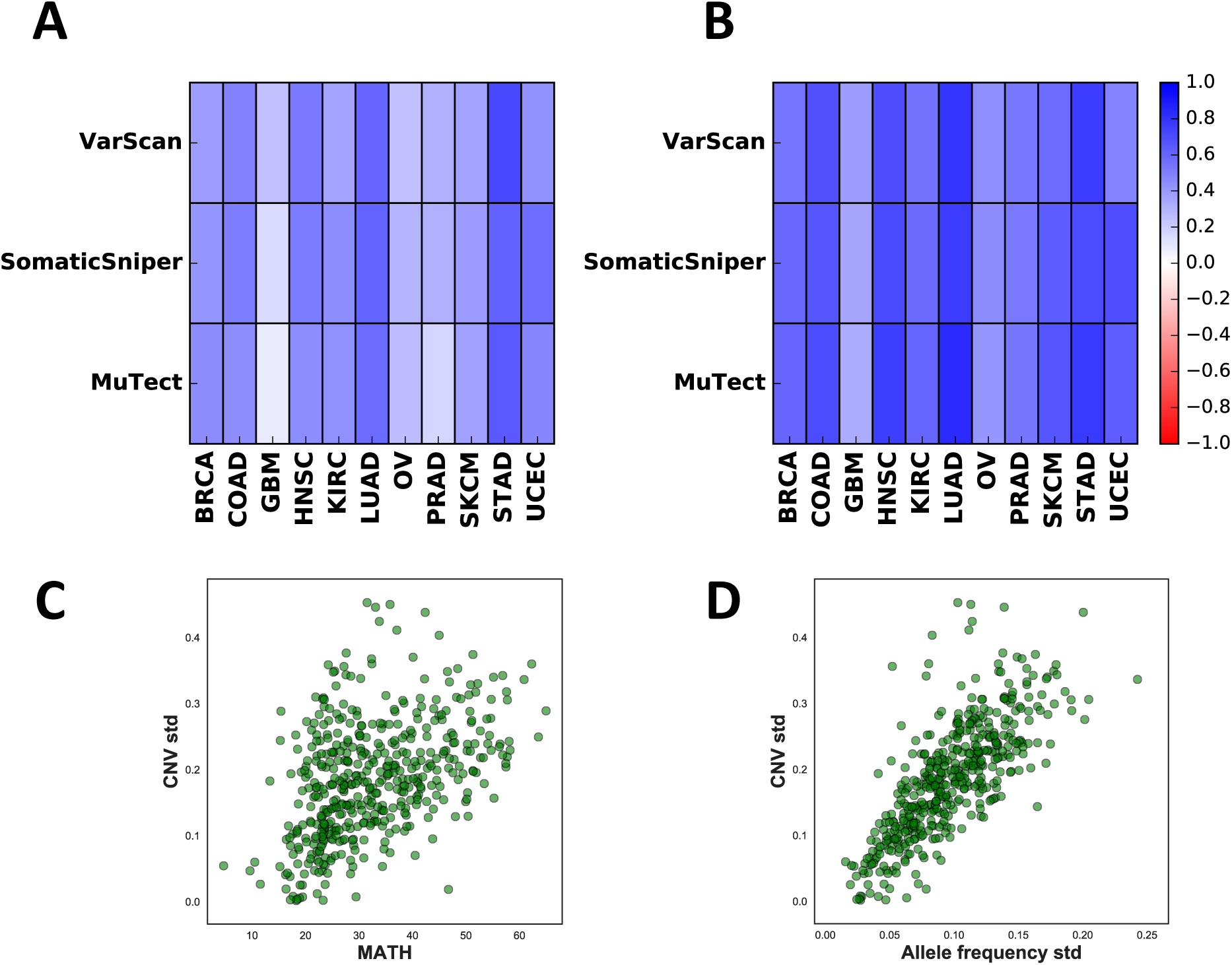
Correlation between allele frequency variations and copy number variations. A) Pearson correlation coefficient of CNV standard deviation and MATH score. B) Pearson correlation coefficient of CNV standard deviation and allele frequency standard deviation. C) CNV standard deviation versus MATH score, plotted for HNSC (mutations called by MuTect). Each circle is an individual sample. D) CNV standard deviation versus allele frequency standard deviation, plotted for HNSC (mutations called by MuTect).

Finally, to check how phenotype is influenced by copy number changes of mutated genes, we analyzed gene expression in UCEC. We compared expression levels of mutated genes having copy number amplification (*CNV* > 1 .5) to the same genes in other tumors lacking mutation and amplification (|*CNV*| < 0.1) (**Figure S12** and *Methods*). We observed that 88% of studied genes exhibited higher expression levels when they had a mutation and were amplified. This suggests that copy number amplification affects survival because it mediates expression changes in mutated genes. We also tested whether this relationship was true when we used CNV standard deviation across the entire genome rather than at just mutated loci. Interestingly, the whole genome value had no predictive power for survival (**Figure S15**). This result indicates that the phenotypic impact arises from amplification of mutated loci, rather than general alteration of copy numbers across the genome.

### Distribution-based measures of intratumoral heterogeneity are consistent with many evolutionary scenarios

The multiple contributions to MATH suggest a theoretical question: what is the uniqueness of MATH score in distinguishing different underlying evolutionary scenarios? To answer this we considered the allele frequency distributions in the linear evolution model [30] and computed MATH score as a function of model parameters, in the simplifying case of no CNVs. The linear model is the simplest model of tumor evolution with selection and assumes that occasional driver mutations lead to selective sweeps in a background of neutral mutations (**Figure 5A**).

**FIG 5.**
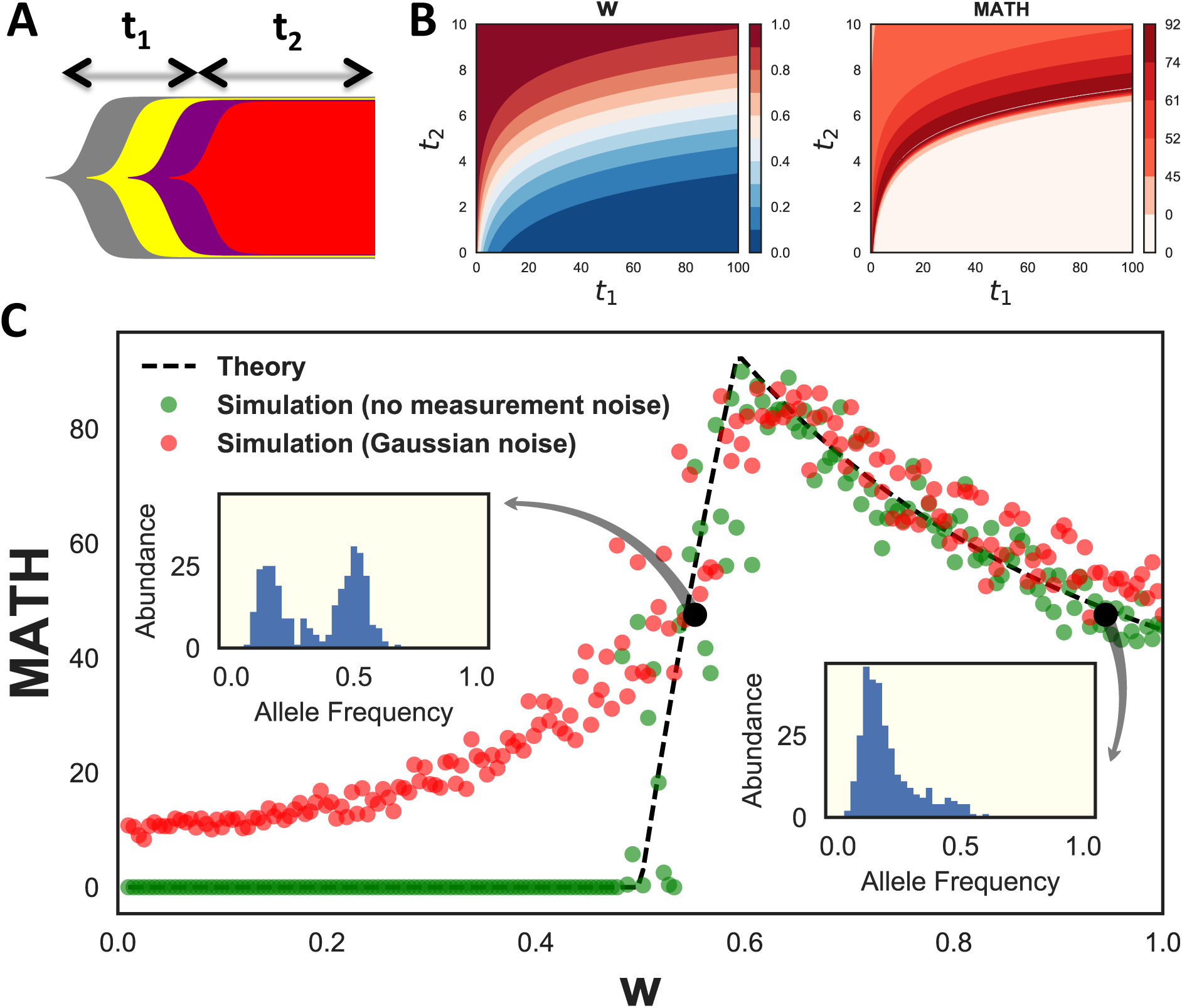
Theoretical analysis of MATH score. A) Schematic diagram of linear tumor evolution model. *t*_1_ is the time from birth of founder cell until the last selective sweep, *t*_2_ is the time from last selective sweep until biopsy. B) Contour plots of neutral fraction, *w*, and MATH as a function of *t*_1_ and *t*_2_ for a noiseless exponentially growing model of linear evolution, *w* is the fraction of total mutations that occurred after the last selective sweep. For this growth model, *w* = 2^*t*_2_^/ (*t*_1_ + 2^*t*_2_^), where time is measured in cell cycles. C) MATH score as a function of neutral fraction, *w*, for a linear tumor evolution model. Dashed lines correspond to analytical calculations. Green dots are samples from the model with 300 mutations. Red dots are simulated results after incorporation of Gaussian measurement noise. Insets correspond to simulated allele frequency distributions for the two black circles, which have equal MATH scores.

The allele frequency distribution can be specified by a variable w that we call the ‘neutral fraction’ (See Methods, equation (4)), i.e. the fraction of total somatic mutations that occurred after the last selective sweep. We analytically derived a closed form for MATH score as a function of *w* (equation (5)). Assuming exponential growth for time-dependence, there are a continuous set of choices of *t*_1_ (the time from birth of the first tumor cell until the last selective sweep) and *t*_2_ (the time since the last selective sweep) that lead to identical values of *w* and MATH (**Figure 5B**). Thus MATH score does not uniquely specify a tumor’s evolutionary history, even in the absence of copy number effects.

To understand the functional dependence of MATH on neutral fraction, we plotted these results and compared them to simulations (**Figure 5C**). These results show two regimes of behavior across *w*. MATH score is zero when *w* ≤ 0.5 (the clonal regime – most mutations are clonal due to the sweep). Addition of noise leads to an increase in MATH score in this clonal regime, but does not significantly affect MATH for *w* > 0.5 (the neutral regime – most mutations have evolved neutrally but vary in when they arose). In the neutral regime MATH is not a one-to-one function of *w*, and two very different allele frequency distributions can have the same score. These results show the limitations of single distribution-based scores to uniquely distinguish tumor evolutionary scenarios. Further features of the distribution, such as bimodality, should be used to resolve this degeneracy.

## DISCUSSION

In a pan-cancer analysis of 4722 samples we explored the robustness of allele frequency distributions to mutation calling and their power for clinical prediction. We demonstrated that the mutation calling process can significantly influence the shape of allele frequency distributions. In general, higher variation in allele frequency was correlated with poorer survival, though this result was statistically significant in only a few cases. The only cancer type with statistically significant results robust across mutation callers was UCEC, but this reflected higher levels of copy number variation along the genome rather than increased subclonal heterogeneity. These results suggest that current single-variable measures of subclonal diversity from exome-seq data are not causally related to clinical outcome.

Our results are consistent with copy number aberrations being a more important predictor of outcome, which has previously been reported over pooled TCGA cancer types [3]. In our analysis most tumor types showed a regular, though not always statistically significant, trend between increased copy number variation and shorter survival time. Notably, high copy number variation has been previously reported to be associated with poor patient survival in serious endometrial carcinoma [31]. When we analyzed TCGA annotated clinical features for associations with CNV standard deviation, we found increased CNV variation for serous carcinoma (**Figure S13**) and high grade tumors.

Ovarian cancer is opposite to the common copy number trend, in that tumors with more CNV variation are associated with longer survival. However, in ovarian cancer BRCA1 deletions often lead to the high-CNV tandem duplicator phenotype, and this phenotype has been found to have better clinical outcome [32]. Indeed, in the TCGA data we observe that increased deletions in BRCA1 are correlated with CNV standard deviation (**Figure S14**), suggesting that the ovarian cancer effect is mediated by the tandem duplicator phenotype.

Despite these findings, we cannot conclude that subclonal heterogeneity is not relevant to survival, as the processes by which resistant populations develop remain poorly understood even though they are crucial to outcome. For example, our work shows that gross estimates of subclonal heterogeneity from exome-seq data have little predictive power, but other studies have shown that resistance can arise from populations too small to be detectable by exome-seq [33]. Higher resolution measurements of subclonal heterogeneity may solve this challenge, and further development of robust computational analyses will be a critical part of such measurements. Clinical data are also still sparse and improving them is a concomitant need.

Finally, we have shown that multiple evolutionary histories can lead to the same allele frequency distributions, and that different allele frequency distributions can have the same MATH score. This degeneracy of scenarios leading to the same measurement is an important reason why new descriptors of heterogeneity are needed. Future surveys over different evolutionary scenarios should be valuable for distinguishing what types of heterogeneity measurements are most likely to reveal features predictive of survival.

## CONCLUSIONS

We studied the robustness of allele frequency distributions to mutation calling procedures across TCGA cancers and explored their prognostic power for patient survival. We found that mutation callers differ significantly in their estimates of the distributions of allele frequencies for individual tumors, and that the association between allele frequency distribution and survival is non-robust to these differences. The major exception is uterine corpus endometrial carcinoma, but the observed effect is mediated by copy number variation rather than subclonal heterogeneity. Our work has implications for cancer precision medicine, as we show that current measures of heterogeneity are not predictive of survival except through an indirect association with copy number variation.

## METHODS

### Computational Time

The average time required to run a task was 2 hours and 25 minutes. We ran a total of 14865 jobs, with total wall time of 1496 days.

### Sample Selection

Samples were chosen only if tumor/normal Illumina exome-seq data existed with alignment to hg19, and corresponding clinical and Affymetrix SNP Array somatic copy number variation data were present. Mutations were called by three mutation callers and samples were discarded if any mutation caller did not find any somatic mutations in that sample. If multiple tumor/normal sample types were available, the smallest sample code was used.

### Mutation Caller Parameters

Mutation calling was done with minor tweaks to the default parameters of each mutation caller (Table I) as shown below.

**TABLE I.** Mutation caller parameters

| Caller | Version | Parameters |
| --- | --- | --- |
| MuTect | 1.1.7 | default |
| SomaticSniper | 1.0.5.0 | -Q 40 -L -G -F |
| VarScan | 2.3.9 | -min-coverage 10<br>-min-coverage-normal 10<br>-min-coverage-tumor 10<br>-min-var-freq 0.05 |

### Postprocessing Filters

We applied post-processing filters on the output VCF files to extract the somatic mutations. For MuTect and VarScan we selected mutations with FILTER=PASS, and in all cases we only used mutations which had zero allele frequency for normal sample and a non-zero value for the tumor. A read depth filter of minimum 50 was also applied to both normal and tumor samples. Furthermore, we discarded mutations which had tumor or normal allele frequency < 0.1. For more detail on mutation calling process and parameters see **Figure S1** and Table I.

### Statistical Distances

To calculate the earth mover distance (EMD)[34], we produced histograms of allele frequencies for each sample using bins of size 0.025 and followed the procedure in [35] for chain-connected spaces. For two probability distributions *f*(*x*) and *g*(*x*) with histogram abundances *f*_*j*_ and *g*_*j*_ and *N* bins, EMD can be written as [35]:

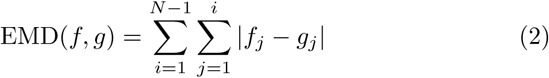

Quantities reported in this paper are the median of this value across each cohort (**Figure 2B**).

We also used another method for quantifying differences between distributions of allele frequencies, where we smoothed the histogram of allele frequencies using Gaussian kernel density estimation with standard deviation *σ* = 0.02, leading to two continuous functions *F*_1_(*f*) and *F*_2_(*f*). We then calculated the cumulative absolute difference of these two functions using 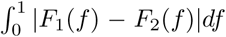. Again the median of cohort was calculated as a single measure of statistical distance (**Figure 2C**).

### Allele Frequency and Read Depth

Read depth and allele frequency were extracted from the FORMAT field of VCF files. Multiallelic and biallelic sites were treated similarly and their allele frequencies corresponded to the ratio of alternative alleles to reference alleles. For MuTect and VarScan the allele frequency computed by the software was directly extracted from VCF files, and for SomaticSniper this was done by using the reference and alternative read counts reported by the software.

### Copy Number Data

For copy number analysis we used copy number level 3 TCGA data measured by Affymetrix SNP Array 6.0 which was already aligned to hg19 and normalized for somatic copy number detection. The quantity used throughout the paper is the *Segment Mean* column in the files which is either used directly or determined at the SNV locus of interest. Wherever mentioned in the text, copy number filtering was done according to [36] where loci with |*CNV*| > 0.2 were removed.

### Copy Number Scaled By Genomic Range

To produce **Figure S15B**, the standard deviation of scaled copy numbers was used. Scaling was done by multiplying the copy number value by the number of bases it covers. This quantity provides a measure of copy number variability which is more influenced by larger copy number events.

### Survival Data

Survival information was gathered from the TCGA data portal by parsing the clinical json files for field *diagnoses.days_to_death* as survival time. In cases where this field was not available, we used *diagnoses.days_to_follow_up* instead. The field *diagnoses.vital_status* was used for determining censored data.

### Survival Analysis

Survival analysis was done using Kaplan-Meier curves and log rank test method from the Lifelines Python package [37]. Only the first 4 years of survival information was used for analyses and any sample with longer overall survival was censored at this time point.

Survival difference values (**Figure S8**) were defined by calculating the Kaplan-Meier curves (**Figure S6**, **Figure S9**) for two groups with high and low variabiltiy score (divided across the median of the score used). We then integrated the absolute value of difference of survival functions:

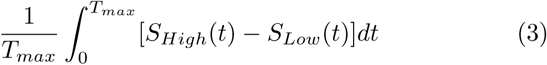

where *S*_*High*_(*t*) and *S*_*Low*_ (*t*) are the survival functions of the two groups and *T*_*max*_ corresponds to the maximum time of data collection (4 years). This definition leads to a quantity between – 1 and 1 which is more negative if the high score group has worse survival, and more positive if the high score group has better survival.

### Expression Analysis

To confirm that copy number amplification influences expression, we downloaded the corresponding RNAseq expression results (RPKM values) of UCEC for each sample from the TCGA database. RPKM values of mutated loci with copy number amplification (defined as *CNV* > 1 .5) from all tumors were selected. We call this quantity *R*_*ij*_, where *i* is the index of sample and *j* is the index of gene. For each value of *R*_*j*_, the RPKM value of gene *j* in the remaining samples was added to a list *S*_*R*_*ij*__ if it was expressed, not mutated, and with no copy number change (defined as |*CNV*| < 0.1). We then averaged the elements of this list to get *S*_*ij*_. This quantity can be thought of as the typical expression level of a gene without any genomic alterations. We then calculated the logarithm RPKM ratio, log(*R*_*ij*_/*S*_*ij*_) as a measure of fold change effect of mutations and copy number amplifications on gene expression. A histogram of this quantity is shown in **Figure S12** where positive values correspond to an increase in expression due to an amplification event.

### Mathematical Modeling

The mathematical model presented in this paper is a linear of model of tumor evolution [30], where any driver mutation leads to a selective sweep and the remaining mutations are passenger mutations. All the mutations before the last selective sweep are clonal with allele frequency 0.5. The remaining mutations follow the neutral model, with cumulative distribution of allele frequency *f* proportional to 1/*f* [19]. Then, the overall allele frequency distribution is the combination of these two distributions sampled according to their occurrence rates (see Supplementary Material):

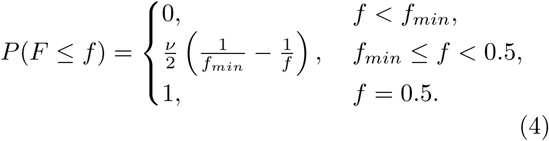

where 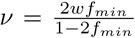, *w* is the ‘neutral fraction’ and is defined as the fraction of all mutations that are neutral, and *f*_*min*_ is the minimum measurable allele frequency. The latter quantity is necessary in order to avoid singularity at *f* = 0. In our simulations we set *f*_min_ = 0.1. The results of simulations with no measurement noise (**Figure 5C**) were calculated using 300 samples from this distribution, and their variations reflect this sampling error.

We derived a closed form for MATH score of the model as a function of *w* (see Supplementary Material for derivation):

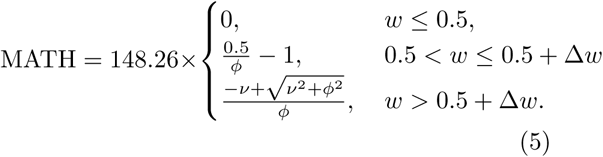

where 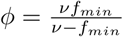, and 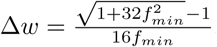.

The allele frequency measurement error can be modeled by a binomial distribution, where observing a mutation on a sequencing read is a Bernoulli trial. For TCGA whole exome sequencing, read depths are quite large (on average 100X) and this error can be approximated by a Gaussian distribution. Therefore, to implement allele frequency noise we drew 300 samples from the distribution in equation (4) and used each resulting value *f* to sample from the normal distribution 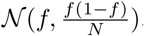. Here *N* = 100 is the sequencing read depth.

### Alignment to GRCh38

To compare our results with alignments to the new genome, we downloaded 227 TCGA breast cancer BAM files of whole exome sequencing data from Cancer Genomics Hub (CGHub) [38]. After converting the bam files to FASTQ files, we remapped the FASTQ files to hg38 reference genome with the Burrows-Wheeler Aligner (BWA) [39]. The somatic mutations were called by MuTect [23]. We found that the allele distributions contained less mutations overall, but their shapes were relatively similar (**Figure S16**)

### Clinical Feature Selection

Random forest feature selction of clinical data (**Figure S13**) was done using *ExtraTreesClassifier* function from Python package scikit-learn [40], with default parameters and 250 estimators. Features with 20% or more unknown values were discarded and the remaining unknown values were set to the median of the feature for numerical features or the mode for categorical features. Data labeling was done by comparing CNV standard deviation to its median.

## ACKNOWLEDGEMENTS

JN would like to thank Joshy George, Francesca Menghi, and Ziming Zhao for helpful comments, Luc Morris for useful discussion and sharing of his previous results, and Anna Lisa Lucido for manuscript edits. JN and SN would like to thank Gaurav Kaushik for instructions on Seven Bridges platform [28]. JHC was supported by the National Cancer Institute of the National Institutes of Health under awards [R21CA191848]

## COMPETING INTERESTS

The authors declare that they have no competing interests.

---

**FIG S1.**
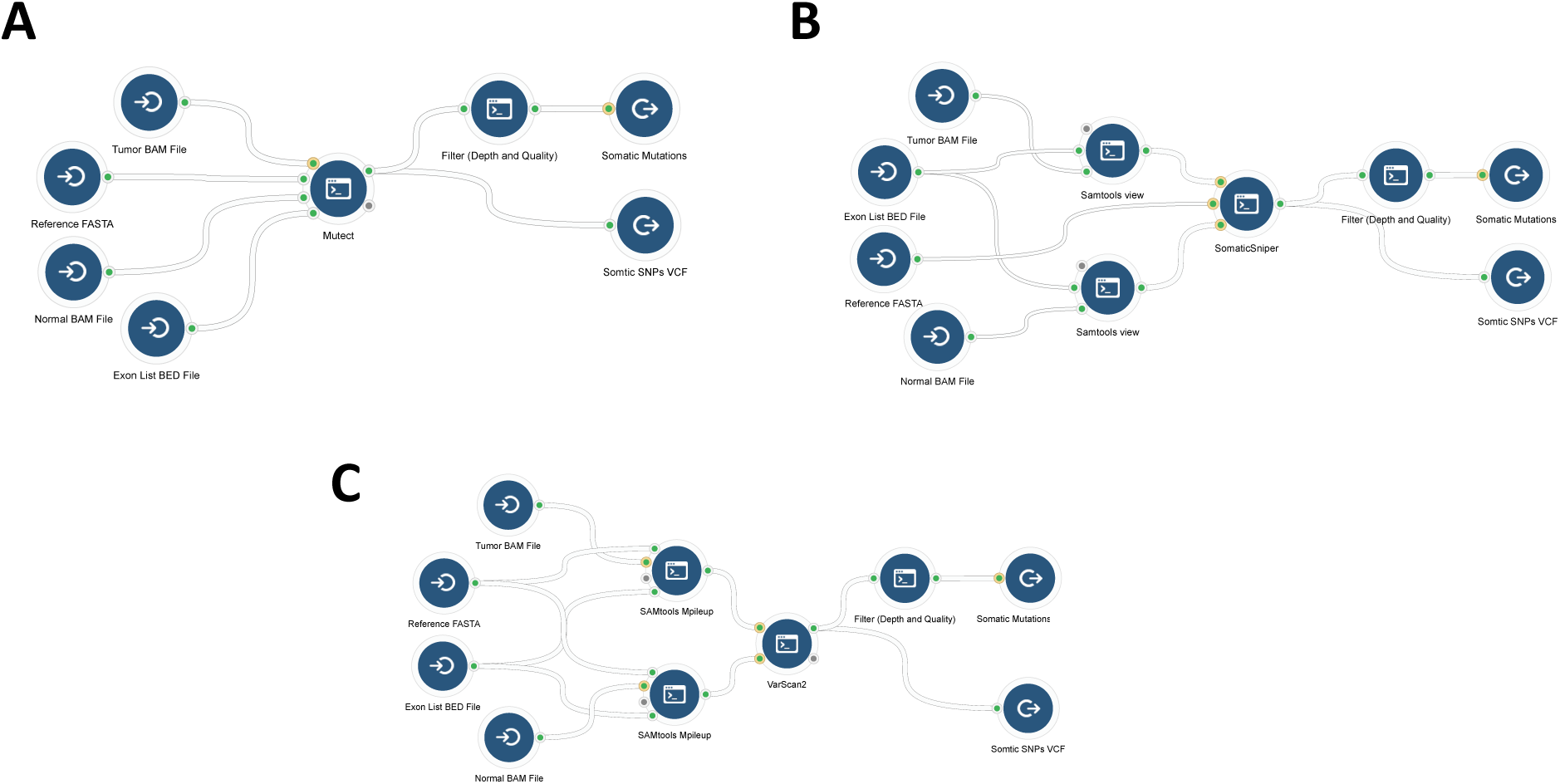
Supplementary Information. Mutation calling pipelines on CGC for A) MuTect, B) SomaticSniper, and C) VarScan

**FIG S2.**
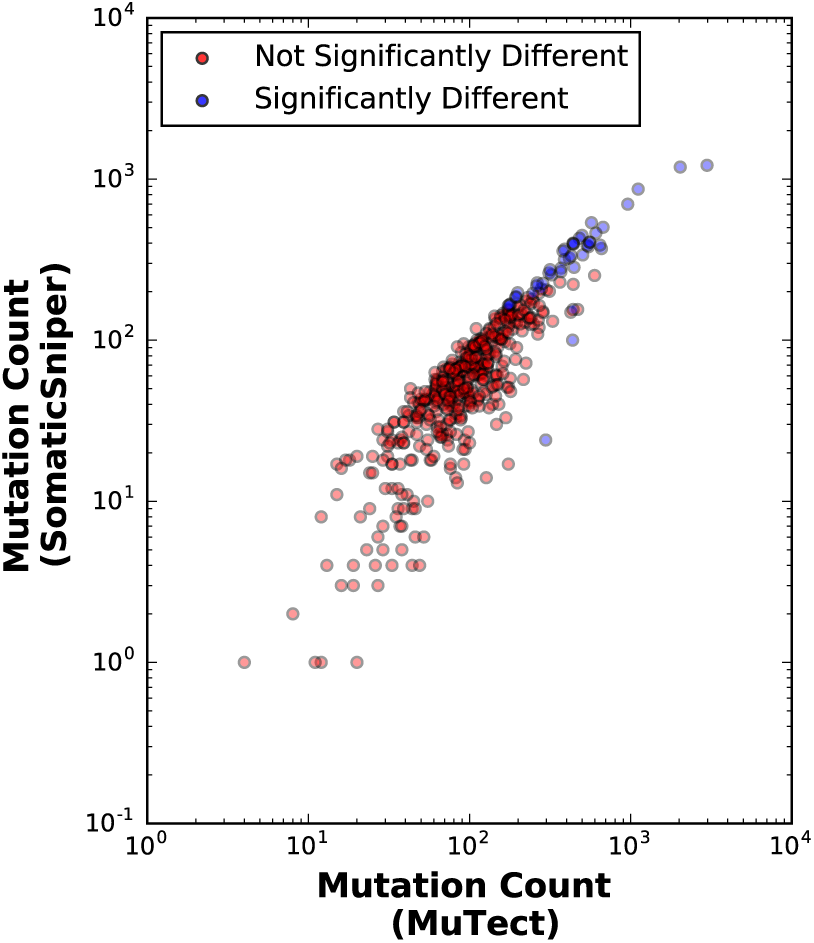
Supplementary Information. Mutation counts for different samples within HNSC compared between two mutation callers. Samples which have distributions that are significantly different according to the Kolmogorov-Smirnov test are shown by blue dots.

**FIG S3.**
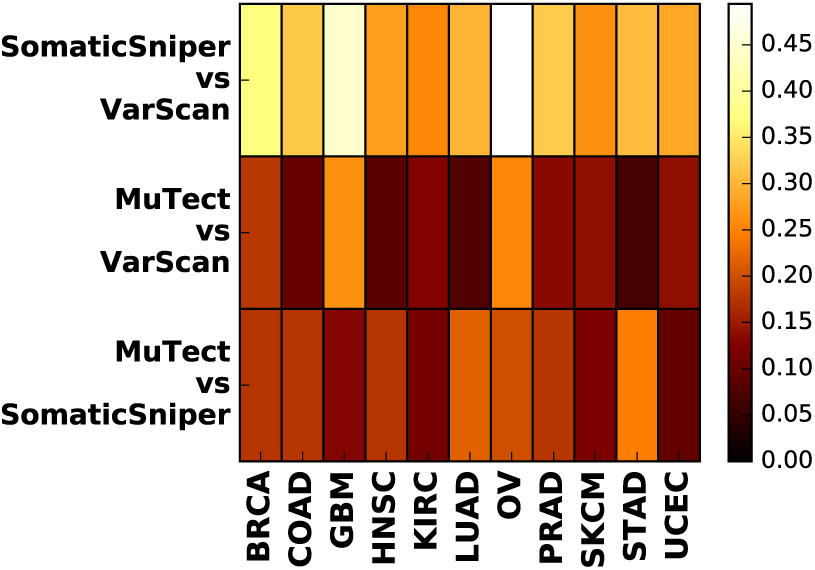
Supplementary Information. Comparison of allele frequency distributions using cumulative absolute difference of smoothed histograms

**FIG S4.**
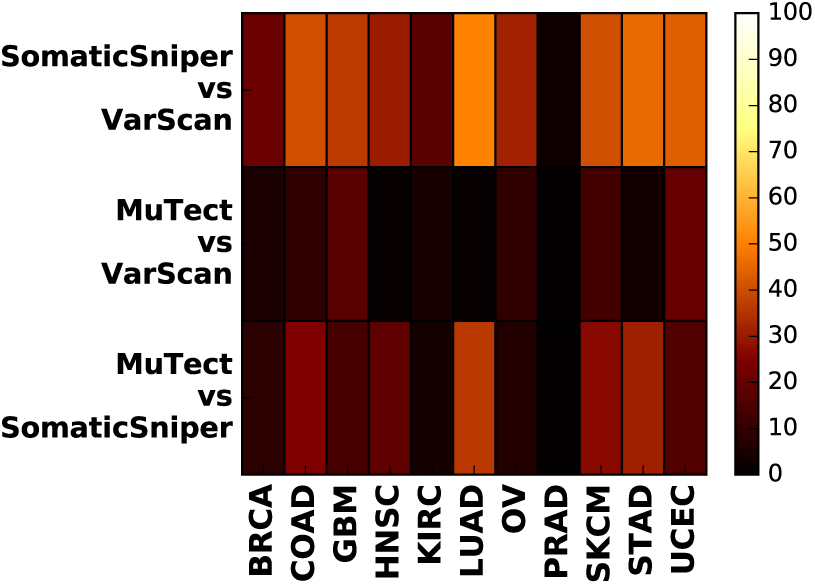
Supplementary Information. Comparisons of allele frequency distributions for different mutation callers. Percentage of significantly different samples (as determined by Kolmogorov-Smirnov test with *p* < 0.05) for mutation caller pairs shown for all cancers with copy number filtering: |*CNV*| < 0.2

**FIG S5.**
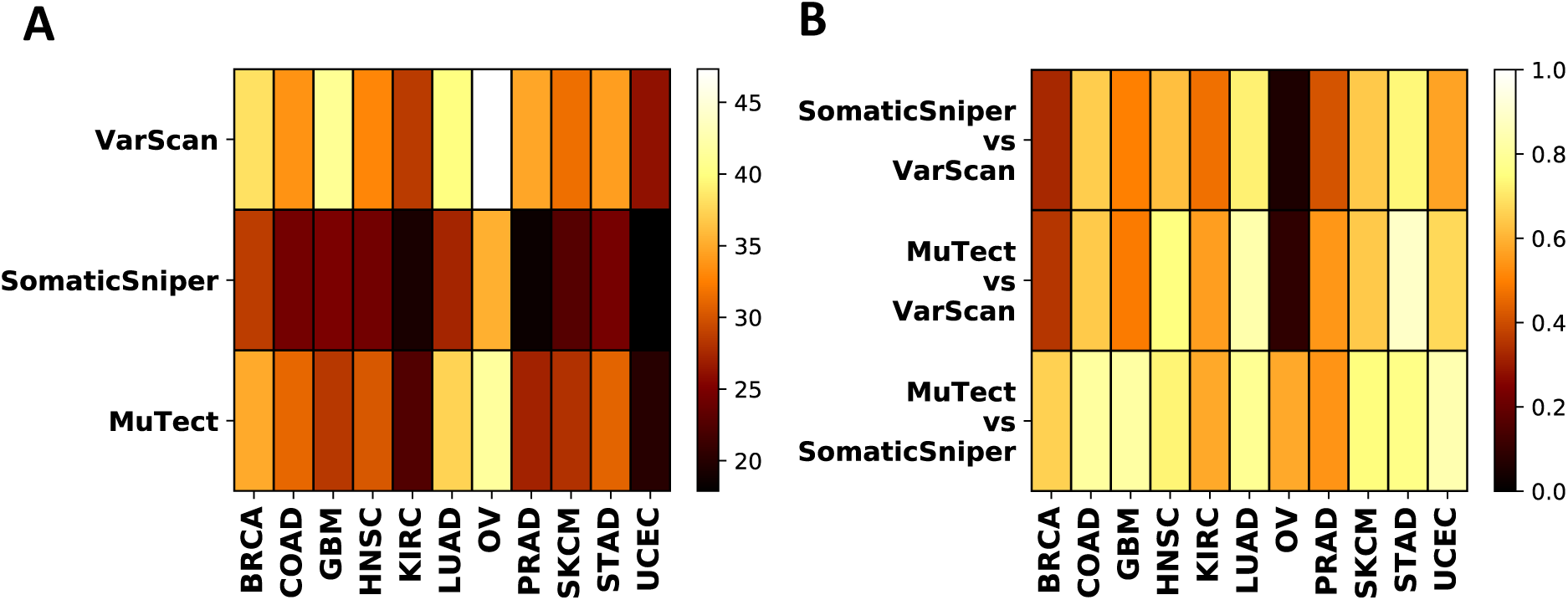
Supplementary Information. A) Median of MATH score for all cancers and mutation callers. B) Spearman correlation coefficient of MATH scores for each cancer type called by pairs of different mutation callers.

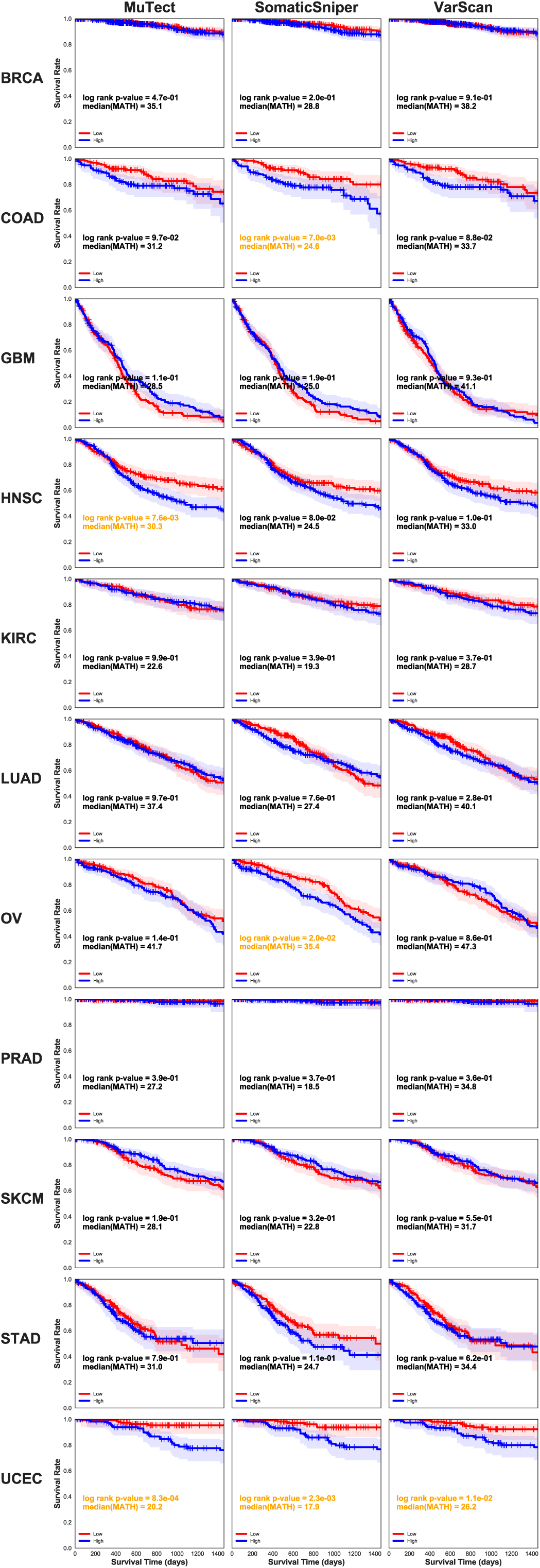

**FIG S7.**
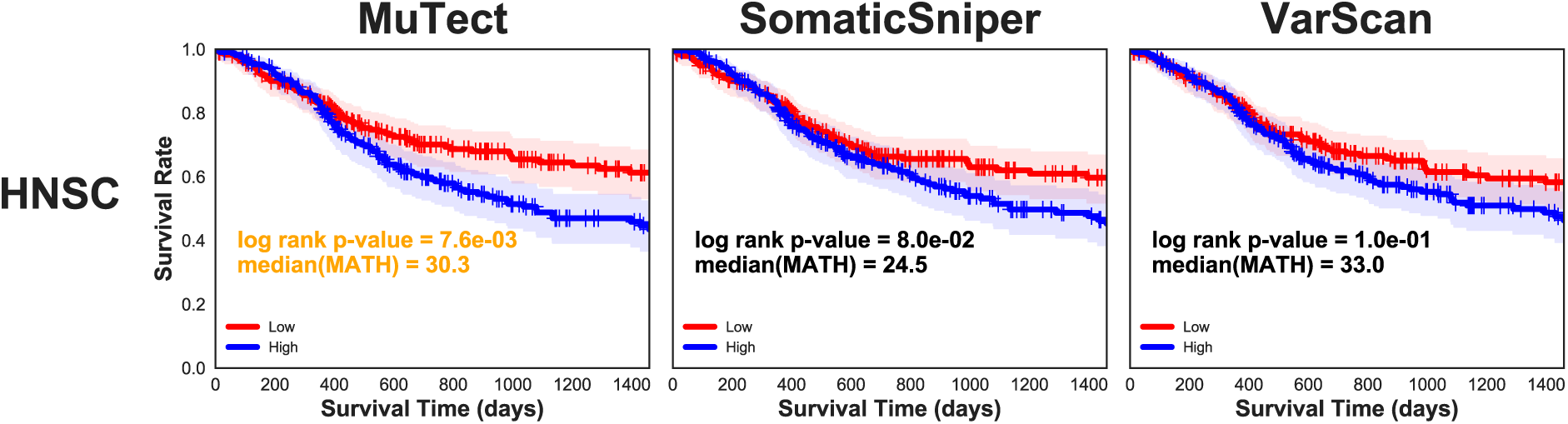
Supplementary Information. Survival analysis using MATH score of the subset of HSNC samples used in [14]

**FIG S8.**
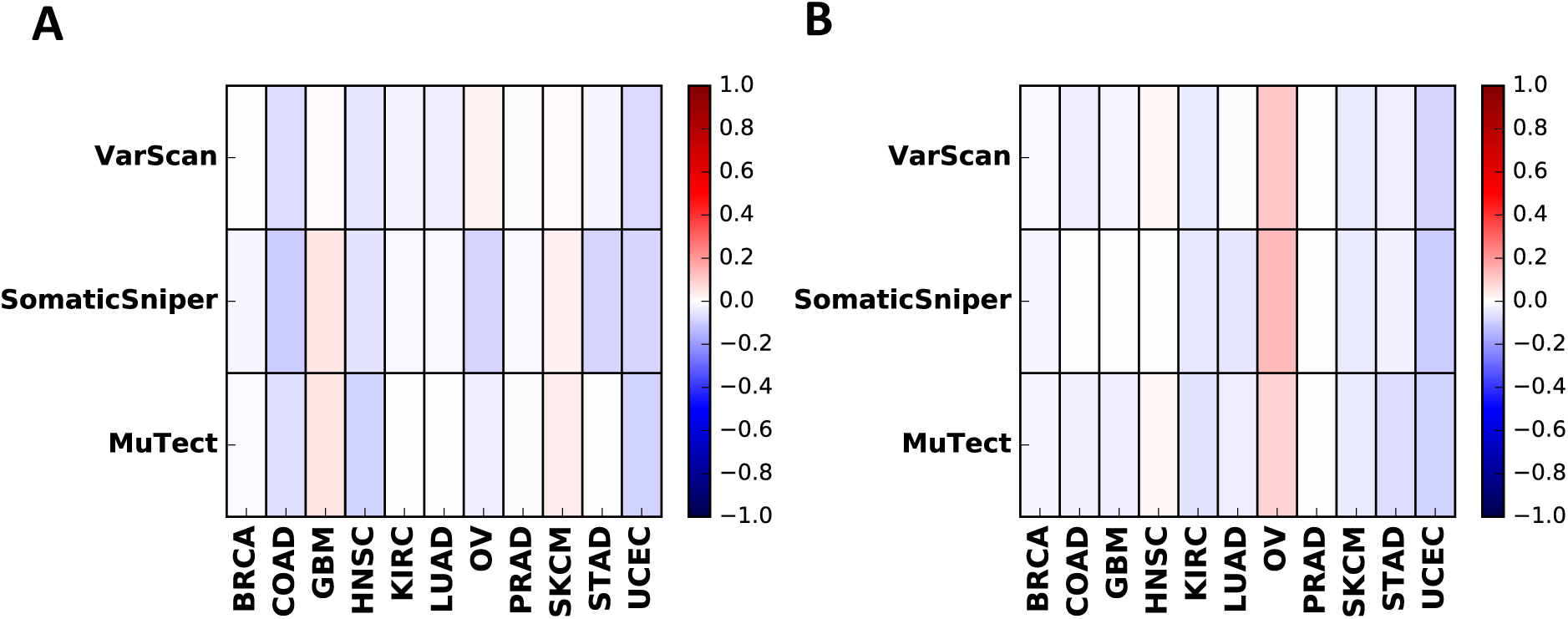
Supplementary Information. A) Survival curve difference for groups separated by MATH score, B) survival curve difference for groups separated by CNV standard deviation

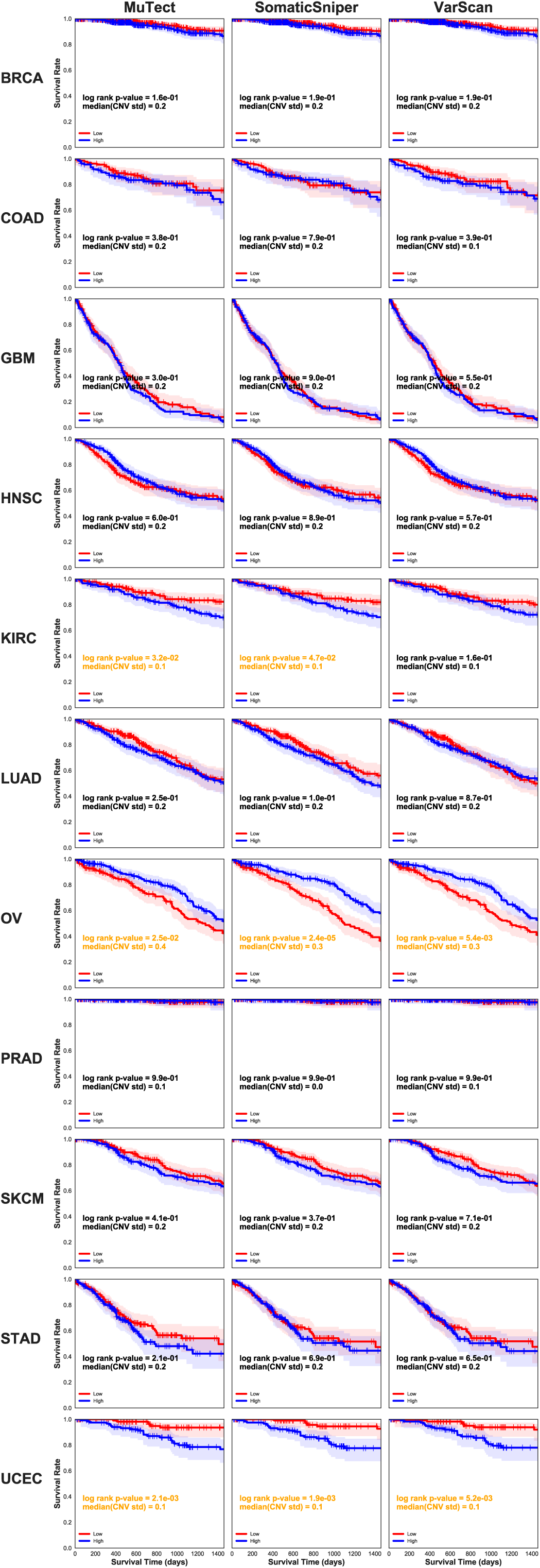

**FIG S10.**
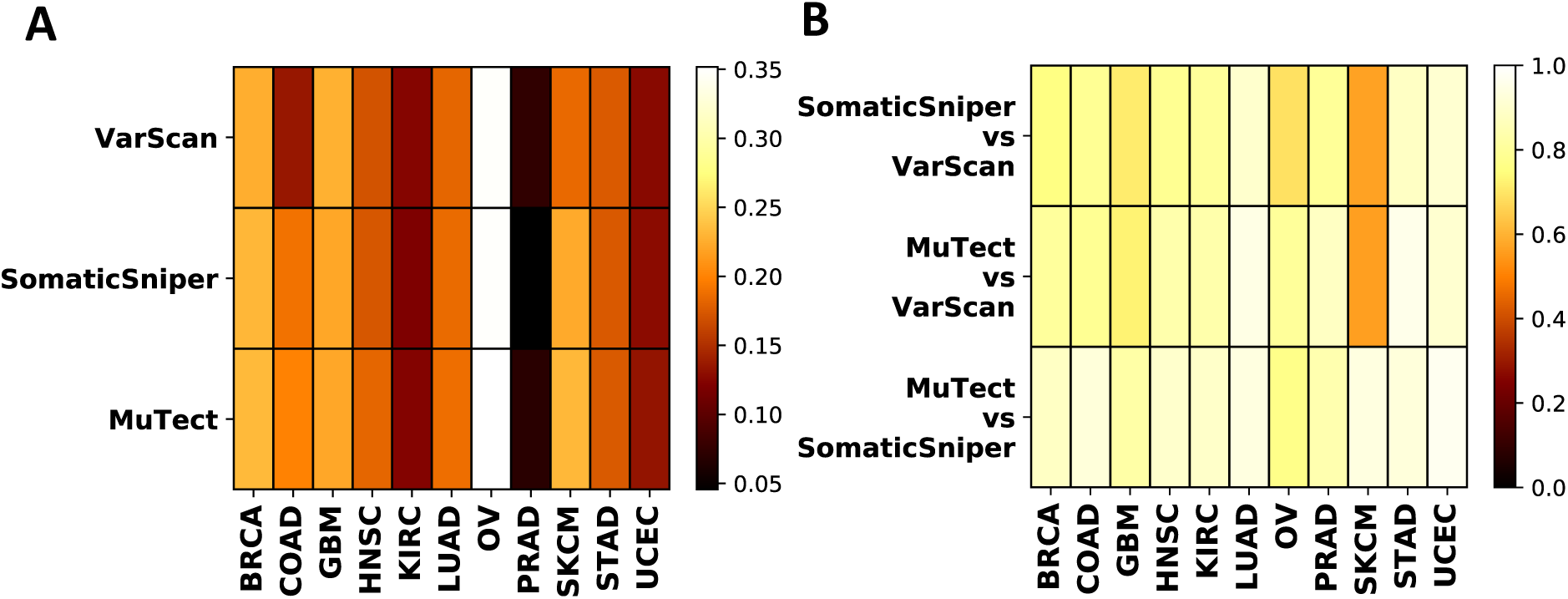
Supplementary Information. A) Median of CNV std score for all cancers and mutation callers. B) Spearman correlation coefficient of CNV std scores for each cancer type called by pairs of different mutation callers.

**FIG S11.**
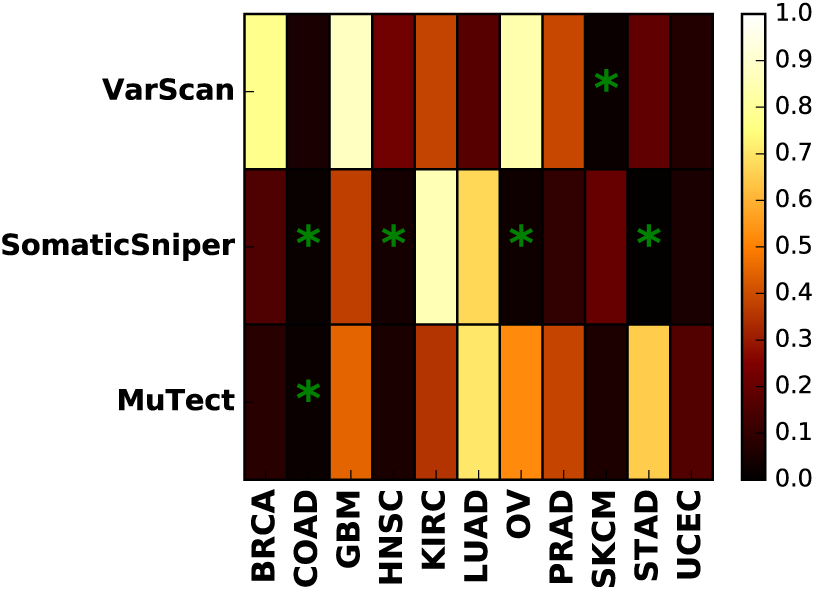
Supplementary Information. Log rank test p-values for comparison of low versus high MATH score when filtered by copy number (|*CNV*| < 0.2). Stars represent significant results as determined by *α* < 0.05

**FIG S12.**
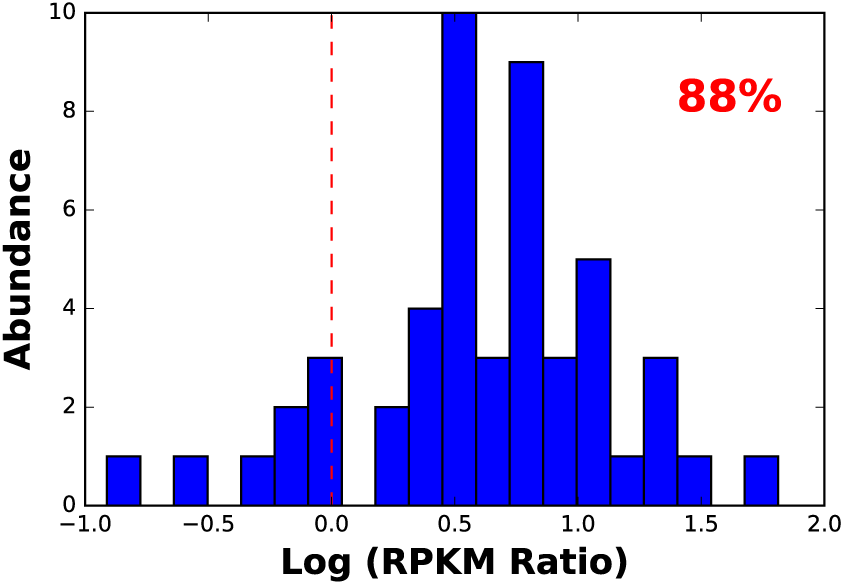
Supplementary Information. Distribution of expression ratio of copy number amplified genes to normal genes for UCEC (refer to *Methods* for detailed definition). This result shows that in 88% of cases copy number amplification leads to an increase in expression of the gene.

**FIG S13.**
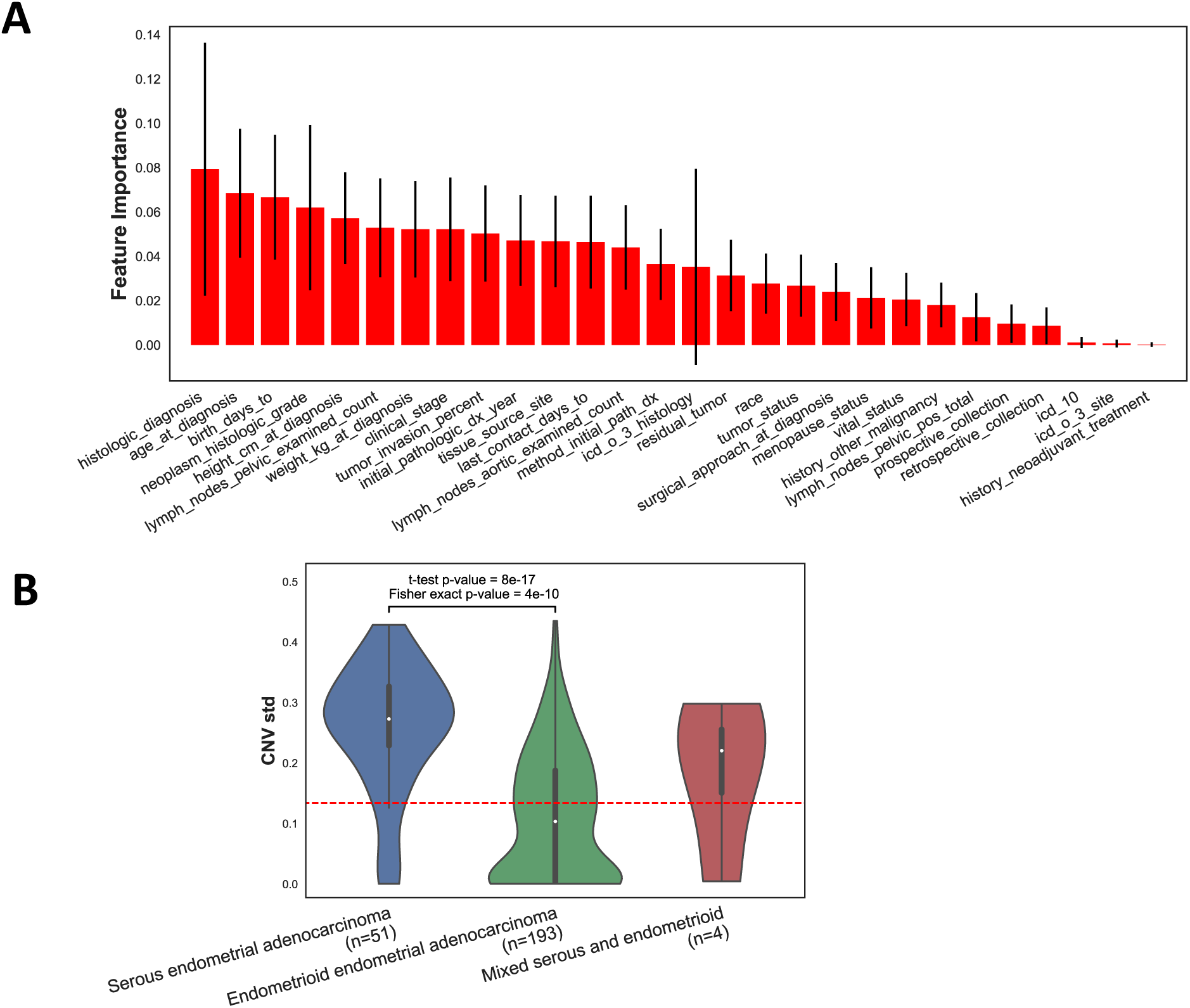
Supplementary Information. A) Clinical data sorted according to their importance in classifying high and low CNV std groups (divided across median) of UCEC. Results were achieved using random forest feature selection. B) Comparison of CNV standard deviation for the three histologic subtypes of UCEC. Dashed red line corresponds to median of CNV standard deviation. The Fisher’s exact test is calculated in comparison to this line.

**FIG S14.**
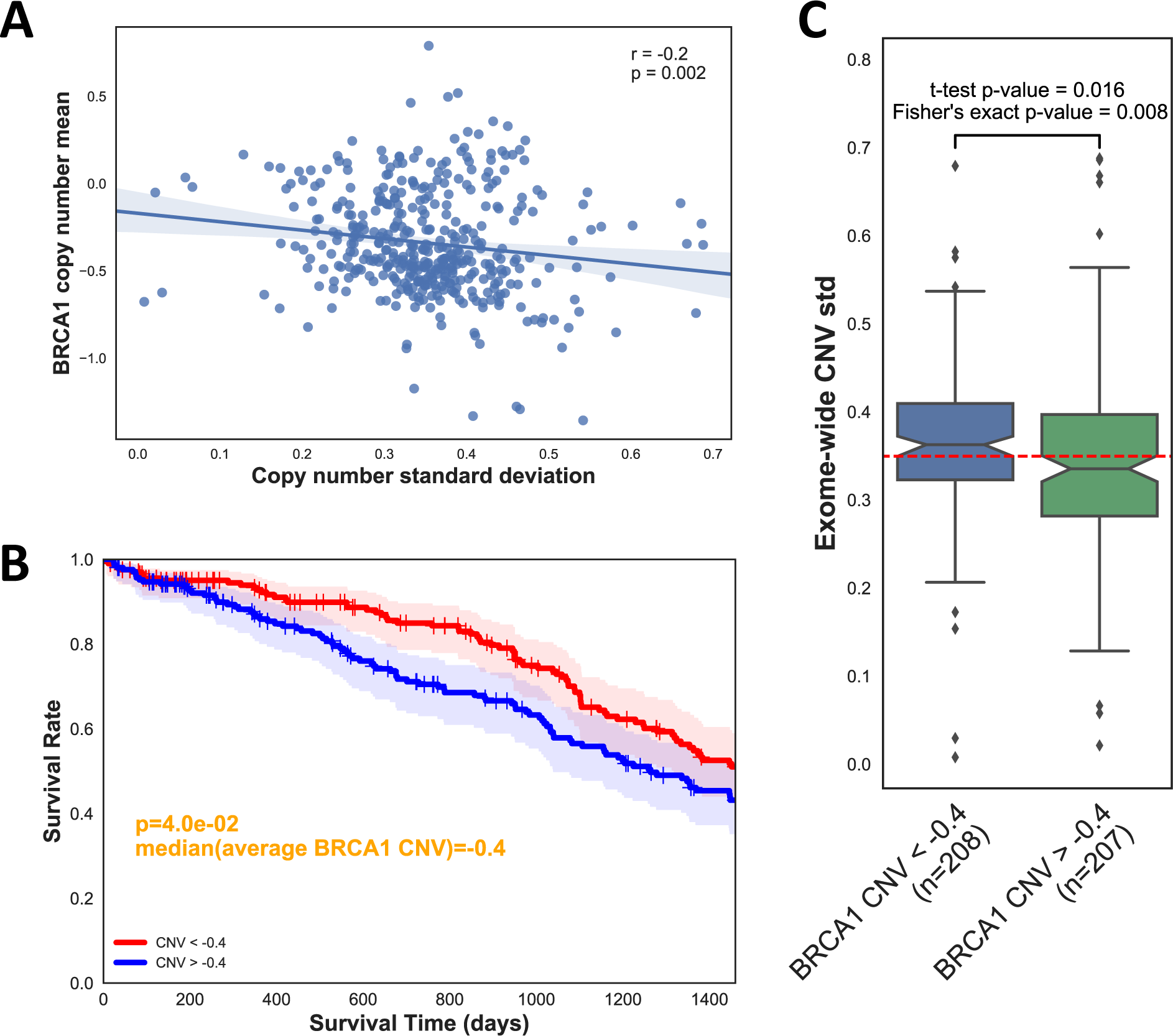
Supplementary Information. Comparison of exome-wide copy number standard deviation (at loci called by SomaticSniper) against BRCA1 copy number for OV. A) Average copy number across BRCA1 plotted against copy number standard deviation showing a negative correlation. Each dot corresponds to one patient. B) Survival analysis of patients divided in relation to median value of BRCA1 copy number average C) Comparison of CNV standard deviation of the two groups in B.Dashed red line corresponds to median of CNV standard deviation. Fisher's exact test is calculated in comparison to this line.

**FIG S15.**
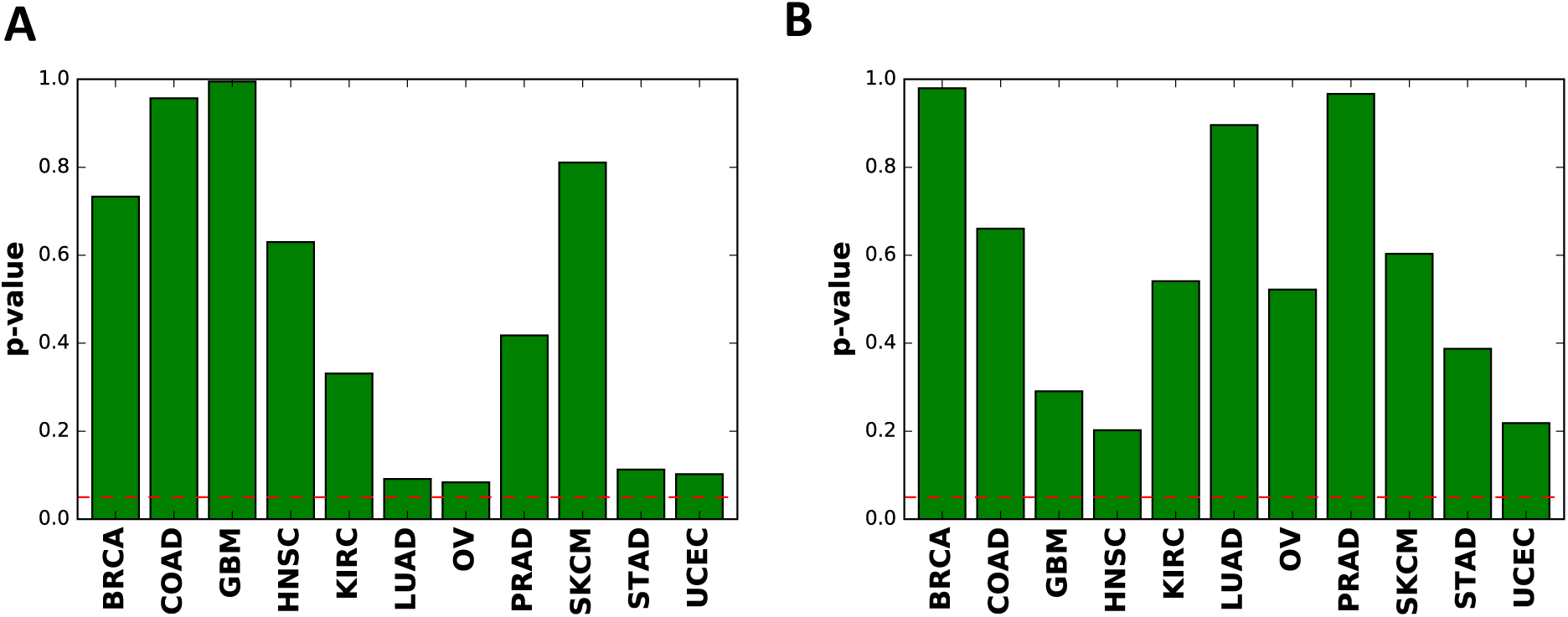
Supplementary Information. Log rank test p-values based on separating patients along median of A) standard deviation of copy number across the genome, or B) standard deviation of copy number across the genome scaled by the corresponding genomic range (see *Methods*). Dashed line corresponds to significance threshold 0.05.

**FIG S16.**
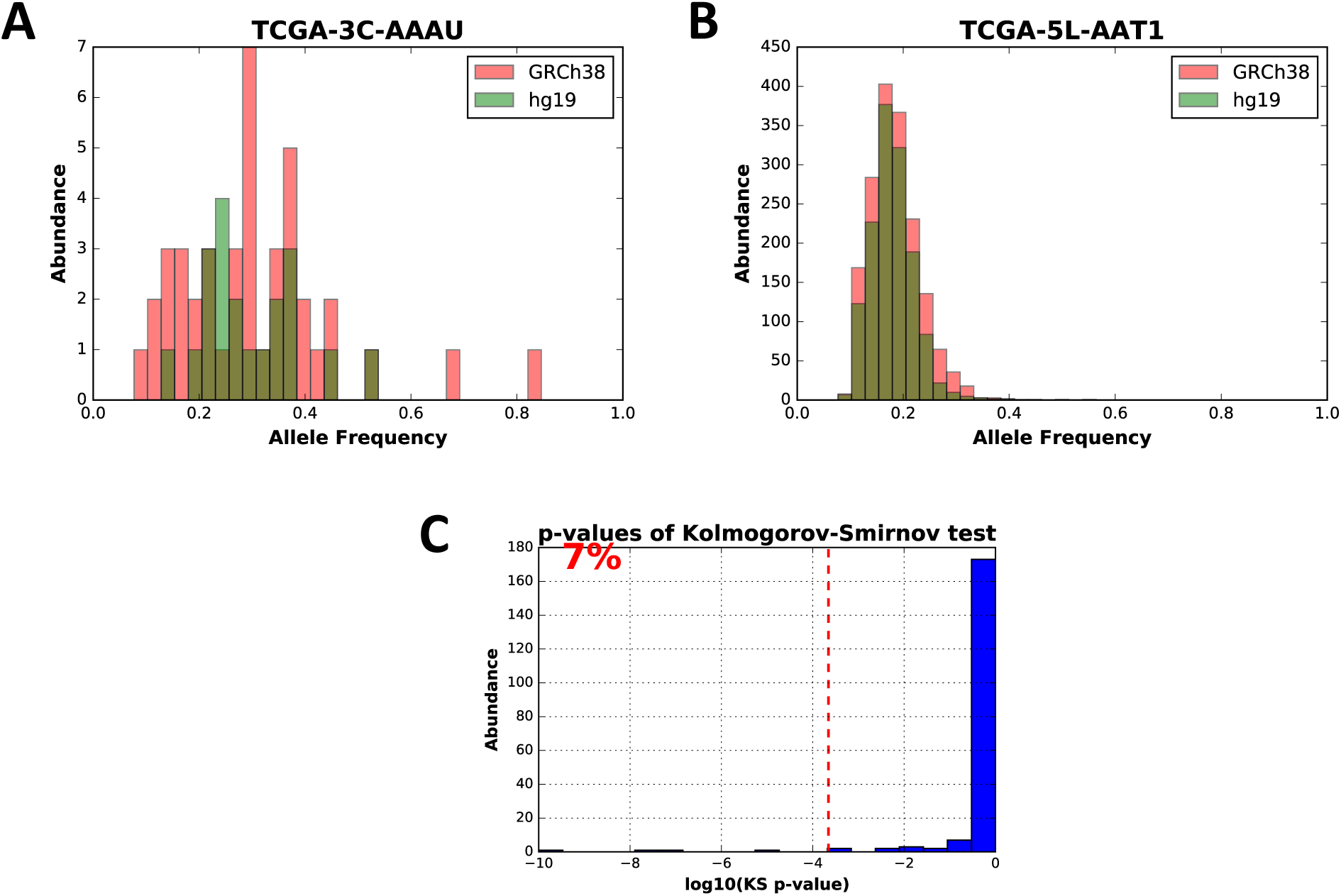
Supplementary Information. A comparison of allele frequency distributions of some BRCA samples aligned to hg19 versus GRCh38. a,b) Two examples of allele frequency distributions. c) Distribution of p-values from the Kolmogorov-Smirnov test between the two alignments. 7% of the samples show significant differences according to Bonferroni corrected significance level 0.05

**FIG S17.**
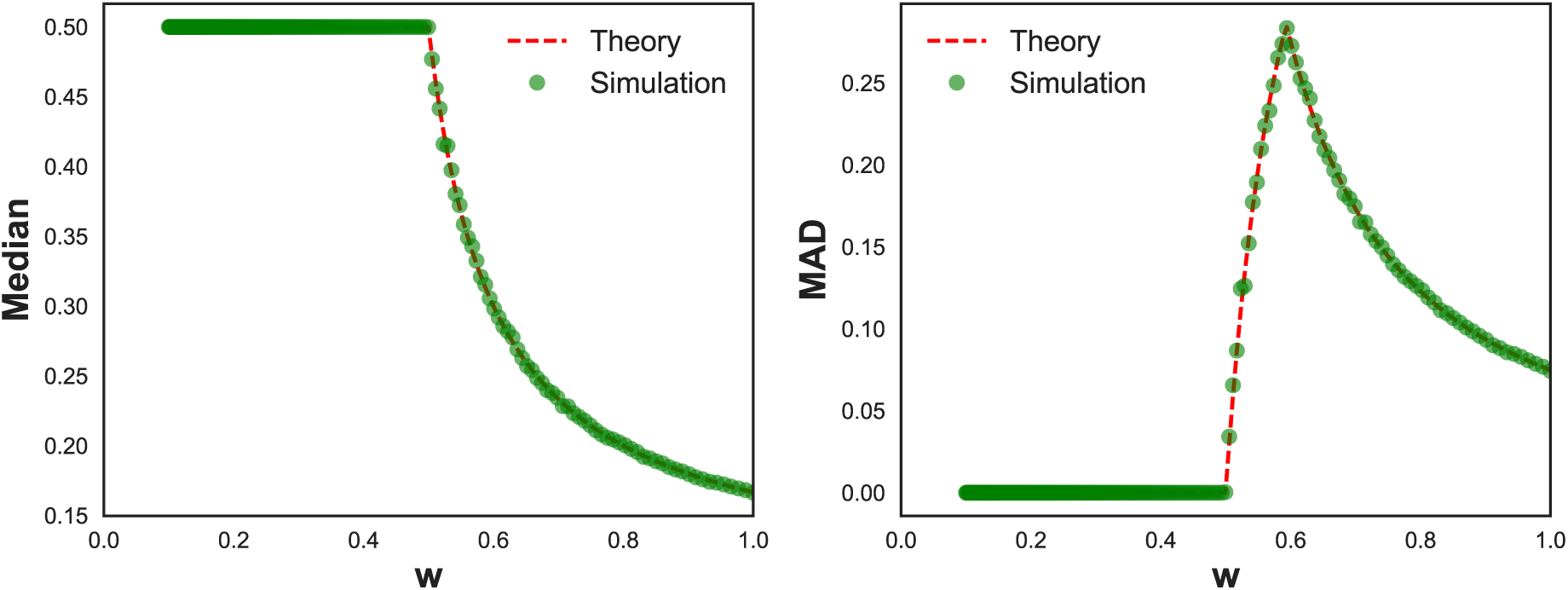
Supplementary Information. Median and median absolute deviation (MAD) of the linear evolution model according to equations (11) and (20). Dashed red line is the theoretical result and green dots are calculated out of 100000 samples from the distribution in equation (8).

## Mathematical Model

The mathematical model described in this paper is based on the linear evolution model of cancer [30]. Most mutations are neutral; however occasionally driver mutations occur which lead to fast selective sweeps. Allele frequencies are determined by the timing of the last selective sweep, and the history of tumor can be divided into two time periods in relation to this event. Consequently, we assume that there are two sets of somatic mutations in the tumor. The subclonal mutations which occurred after the last selective sweep, and the clonal mutations which occurred before the last selective sweep. The fraction of mutations in these two groups will be represented by *w* and 1 – *w* respectively. The distribution of neutral subclonal mutations follows [19]:

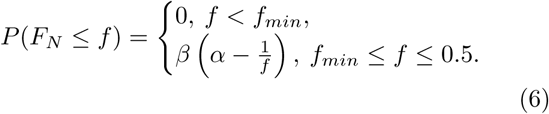

where *F*_*N*_ is the random variable associated with allele frequencies of the neutral model. In order to have *P*(*F*_*N*_ ≤ *f*_*min*_) = 0 and *P*(*F*_*N*_ ≤ 0.5) = 1, we set 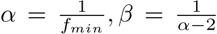 where *f*_*min*_ is the minimum allele frequency measured. All the clonal mutations have allele frequency 0.5 and their distribution is:

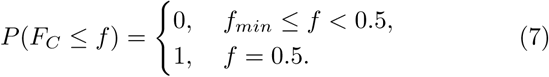

where *F*_*C*_ is the random variable associated with clonal mutations. For allele frequency *F* = *F*_*N*_ ∪ *F*_*C*_, the overall probability distribution will be sum of probabilities (given that *F*_*N*_ ∩ *F*_*C*_=0) weighted by their rate of occurrence:

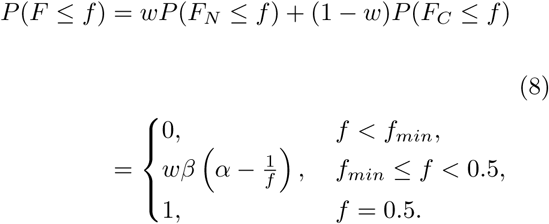

To calculate MATH score for this distribution, we will calculate the median and median absolute deviation (MAD) separately.

### Median

We will denote the median of allele frequencies by *ϕ*. In general *ϕ* has to satisfy the following inequalities:

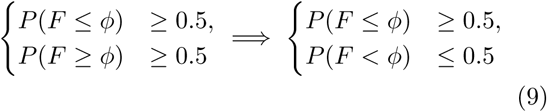

For *ϕ* = 0.5 the first row of equation (9) is trivial (1 ≥ 0.5), and the second row leads to *w* ≤ 0.5. On the other hand, for *f*_*min*_ ≤ *ϕ* < 0.5, equation (9) reduces to:

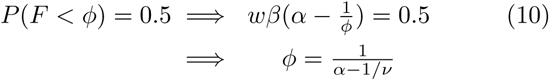

where *ν* = 2*wβ*, which leads to *ϕ* < 0.5 only if *w* > 0.5. To summarize our results, the median of allele frequencies can be written as:

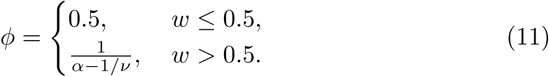

Sampling from the distribution in equation (8) confirms this result (**Figure S17**).

### Median Absolute Deviation (MAD)

We will denote MAD by *m*. It can be derived from the following relationships:

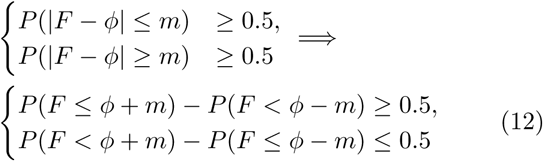

If *w* ≤ 0.5 we have *ϕ* = 0.5. We start by assuming that *m* > 0. In this case *ϕ* + *m* > 0.5, leading to *P*(*F* ≤ *ϕ* + *m*) = 1 and *P*(*F* ≤ *ϕ* + *m*) = 1. Hence equation (12) can be simplified to:

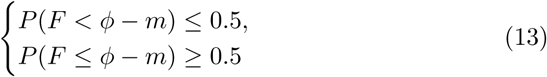

Since *ϕ* – *m* < 0.5 the functions are continuous and we can instead write *P*(*F* ≤ *ϕ* – *m*) = 0.5. But *P*(*F* ≤ *ϕ* – *m*) ≤ *P*(*F* < 0.5) < *w* which cannot be true, given that *w* ≤ 0.5. As a result *m* cannot be positive. Since *m* is non-negative we conclude that *m* = 0.

On the other hand, for *w* > 0:5, we have 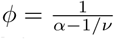. this case there are four possibilities for solving equation (12) that we will separately explore:

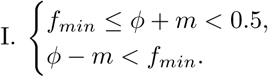

Functions are continuous in this region. So *P*(*F* ≤ *ϕ*+*m*) = 0:5, which can be solved similarly to equation (11) and gives *ϕ*+*m* = *ϕ* ⇒ *m* = 0. However this cannot be true, because the assumptions of this case can only be true for positive *m*.

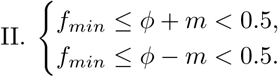

Functions are continuous in this region and we can write:

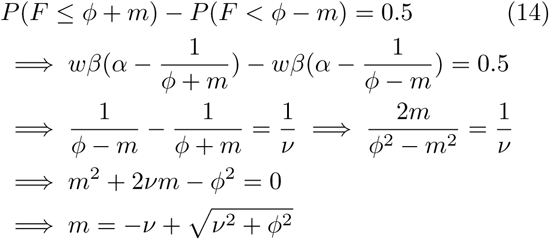

This result is bounded by *m* = 0:5 – *ϕ*.

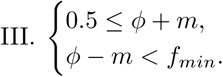

Functions are not continuous in this region. We can write the second row of equation (12) as

*P*(*F* < *ϕ* + *m*) ≤ 0:5. But *P*(*F* < *ϕ* + *m*) ≥ *P*(*F* < 0:5) = *w* > 0:5. Hence this cannot be true.

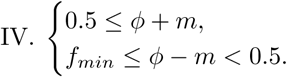

Functions are not continuous in this region. We can write the first row of equation (12) as:

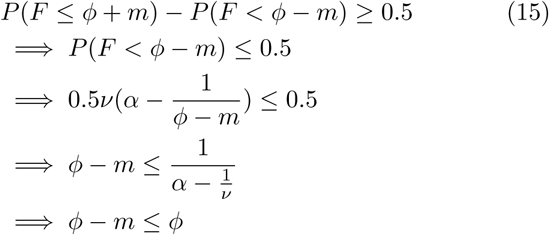

which is a trivial result. For the second row of equation (12) we have:

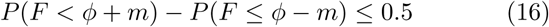

If *ϕ* + *m* > 0:5 equation (16) leads to:

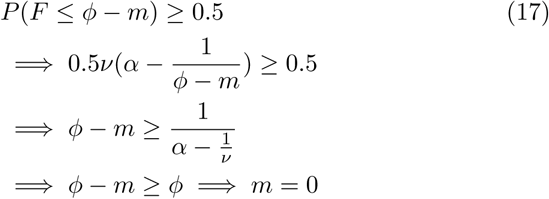

which does not satisfy the assumptions of this case and cannot be true. On the other hand, if *ϕ* + *m* = 0:5 equation (16) is equal to:

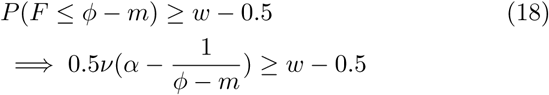

After some calculation we find:

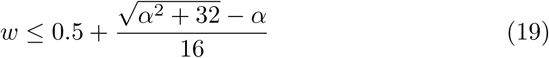

In conclusion, only cases II and IV lead to acceptable solutions. To summarize these results, for MAD we have:

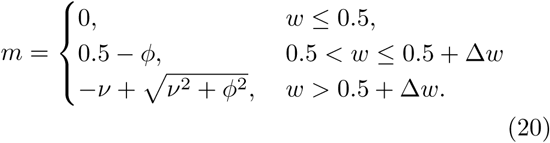

where 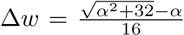. Sampling from the distribution in equation (8) confirms this result (Figure S17).

### MATH Score

MATH can be derived by the following formula:

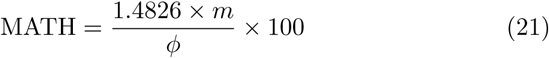

where the constant is the scale factor for median absolute deviation. Using equations (11) and (20) we can write this as:

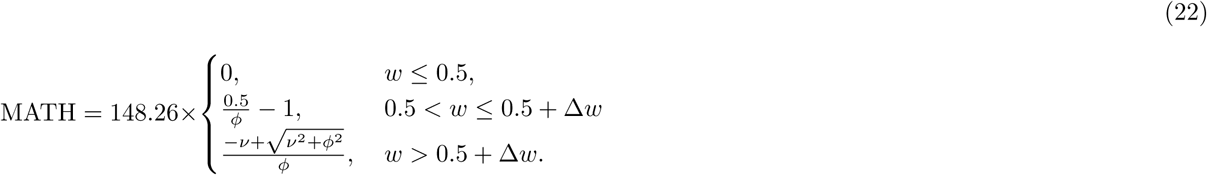

